# Genomic Analyses Suggest No Risk of Vancomycin Resistance Transfer by Strain VE202-06

**DOI:** 10.1101/2024.09.18.613467

**Authors:** Andrea R. Watson, Wei-An Chen, Willem van Schaik, Jason M. Norman

**Author notes:** Corresponding author: Jason M. Norman, For general inquires.

## Abstract

In 2016, the US Food and Drug Administration published guidance for the early development of live biotherapeutic products (LBPs)^1^. Of particular importance is the characterization of LBP strains and the potential transfer of antimicrobial resistance (AMR) genes to relevant microbial organisms in the recipient’s microbiota. Van der Lelie et al^2^, make unsupported claims that the LBP strain VE202-06 encodes a transferable vancomycin resistance element. Here we provide our analysis of the potential transfer of AMR by strain VE202-06. These data indicate that strain VE202-06 has no risk of transferring AMR to relevant microbial organisms.

## INTRODUCTION

Modulation of the microbiome by donor-derived procedures (i.e. fecal microbiota transplant [FMT]) and LBPs (i.e. defined consortia) have proven effective in treating human gastrointestinal infections and disease. Vedanta Biosciences currently has 2 active, international clinical programs to evaluate defined bacterial consortia. The RESTORATiVE303 study (NCT06237452) is a Phase 3 study to evaluate the safety and efficacy of an 8-strain defined consortium, VE303, for the prevention of recurrent *Clostridioides difficile* infection (rCDI). The COLLECTiVE202 study (NCT05370885) is a Phase 2 study to determine the safety and efficacy of a 16-strain defined consortium, VE202, in patients with mild-to-moderate ulcerative colitis (UC). Prior to initiation of these clinical trials, nonclinical and clinical safety data were provided to, and evaluated by, multiple international regulatory authorities. The nonclinical data included genomic characterization of the potential for AMR gene transfer from LBP strains to resident bacterial strains. Van der Lelie et al^2^ specifically reference the presence of transferable vancomycin resistance elements in the genome of a VE202 consortium strain, *Blautia coccoides* (VE202-06; GenBank Accession Number Accession: PRJDB525), without providing data to support their statement. Herein, we provide evidence that refutes that statement, to underscore the steps taken to avoid the risk of transferrable resistance in the rational design of LBPs, before they enter the clinic.

Vancomycin is used routinely for the treatment of Gram-positive bacterial infections and resistance is of great concern, particularly in *Enterococcus* species. Vancomycin resistance is mediated through the replacement of the terminal D-Ala-D-Ala moiety in peptidoglycan peptide stems by D-Ala-D-lactate (D-Lac), or, more rarely, by D-Ala-D-serine (D-Ser). Vancomycin binds to D-Ala-D-Ala with high affinity, preventing cross-linking of peptidoglycan stem peptides by penicillin-binding proteins, leading to cell death. The affinity of vancomycin to D-Ala-D-Lac or D-Ala-D-Ser is >1000- and 7-fold lower, respectively, thus conferring resistance^3^.

Current understanding of how microbes can become resistant to vancomycin indicates that there are two general mechanisms: 1) acquisition of genes from other bacteria in the community via horizontal gene transfer (HGT); or 2) presence of native genes that encode the biochemical machinery that leads to a different terminal residue than D-Ala in peptidoglycan stems. This latter route to resistance explains why many lactobacilli and other Gram-positive bacteria are intrinsically resistant to vancomycin^4,5^. The statement “the presence of transferable vancomycin resistance elements as found in the genome of the VE202 consortium strain *Blautia coccoides*”^2^ can only be interpreted as referring to the potential for HGT of vancomycin resistance from strain VE202-06. Transferable vancomycin resistance has been intensively studied in *Enterococcus* and has led to the identification of multi-gene operons that catalyze the remodeling of the terminal residue of the peptide stems of peptidoglycan^6^. These operons are frequently associated with integrative and conjugative elements, and plasmids and can thus spread via HGT^7^.

## RESULTS AND DISCUSSION

To determine the presence of vancomycin resistance genes and their proximity to known mobile genetic elements (MGEs), we have conducted analyses using two independent workflows. In Workflow 1, the genomes were annotated using the National Center for Biotechnology Information’s (NCBI) Prokaryotic Genome Annotation Pipeline (PGAP)^8^. In Workflow 2, the genomes were annotated by Bakta^9^ v1.4.0 database v3.1 (AMR, virulence genes, and MGEs), Abricate^10^ v1.0.1 (MGEs), ViralVerify^11^ v1.1 (plasmids and viruses), and geNomad^12^ v1.5.2 (plasmids and viruses). Because the annotation of bacterial genomes and assessment of synteny is influenced by genome quality and completeness, we performed our analysis on two strain VE202-06 assemblies with different levels of fragmentation. (**Table 1**). The analysis of the more fragmented genome was necessary to compare our results with the published statement.

**Table 1.** Summary of AMR Gene Predictions in Strain VE202-06.

|  | 2013 Assembly | 2022 WCB Assembly |
| --- | --- | --- |
| GenBank Accession | GCA_000508925.1 | CP166247.1 |
| Sequencing technology | Ion PGM, 454 FLX | Oxford Nanopore GridION, Illumina NextSeq 2000 |
| Assembly method | Newbler v. 2.8 | Flye v2.9<br>Polished with Medaka v1.4.4,<br>PolyPolish v0.5.0, and MaSuRCA (POLCA) v.4.0.9 |
| Genome size | 6 Mb | 6 Mb |
| Contigs (N) | 327 | 1 [circular] |
| Contig N50 | 38.7 kb | 6 Mb |
| Genome coverage | 28.0x | 95x |
| **Workflow 1: PGAP v.2024-07-18.build7555** |  |  |
| Predicted AMR gene | VanZ<sup>a</sup>; contig=BAHT02000064.1; start=12,138,<br>VanR; contig=BAHT02000086.1; start=8,958,<br>VanZ<sup>a</sup>; contig=BAHT02000139.1; start=13,434,<br>VanZ<sup>a</sup>; contig=BAHT02000166.1; start=16,904,<br>VanZ<sup>a</sup>; contig=BAHT02000202.1; start=10,621,<br>VanZ<sup>a</sup>; contig=BAHT02000202.1; start=18,634 | VanZ<sup>a</sup>; start=154,194,<br>VanZ<sup>a</sup>; start=162,207,<br>VanZ<sup>a</sup>; start=3,214,988,<br>VanR; start=3,640,511,<br>VanZ<sup>a</sup>; start=4,591,356,<br>VanZ<sup>a</sup>; start=5,478,254,<br>VanY; start=6,158,912 |
| **Workflow 2: AMRFinderPlus, Bakta (UniRef BLAST & AMRFinderPlus), PlasmidFinder, ICEberg2, geNomad, ViralVerify** |  |  |
| Predicted AMR Gene | VanZ<sup>a</sup>; contig=BAHT02000064.1; start=12,138 [chromosome],<br>VanR; contig=BAHT02000086.1; start=8,958 [uncertain],<br>VanZ<sup>a</sup>; contig=BAHT02000139.1; start=13,434 [chromosome],<br>VanZ<sup>a</sup>; contig=BAHT02000166.1; start=16,904 [chromosome],<br>VanZ<sup>a</sup>; contig=BAHT02000202.1; start=10,621 [plasmid] | VanZ<sup>a</sup>; start= 154,194 [chromosome],<br>VanZ<sup>a</sup>; start=3,214,988 [chromosome],<br>VanR; start= 3,640,511 [chromosome],<br>VanZ<sup>a</sup>; start=4,591,356 [chromosome],<br>VanZ<sup>a</sup>; start=5,478,254 [chromosome] |

<sup>a</sup> VanZ family and VanZ domain-containing proteins

Workflow 1, using the default PGAP parameters, predicted a *vanR* gene and several VanZ family or VanZ domain-containing proteins in both VE202-06 assemblies, and a *vanY* gene in the updated, circular genome (**Table 1**). Workflow 2 also predicted *vanR* genes, as well as several VanZ domain-containing and VanZ family proteins, in both VE202-06 assemblies. In the more fragmented VE202-06 assembly, workflow 2 was unable to predict the topology of the region in which *vanR* was found and marked one possible VanZ family protein as plasmid-associated. No plasmids were assembled in the updated, circular VE202-06 genome, thus confirming all *van* genes were located on the chromosome, including the previously plasmid-associated VanZ family protein (**Figure 1**). Importantly, no complete multi-gene *van* operons were predicted in either genome by either workflow, and no *van* genes were associated with known MGEs (**Figure 1** and **Supplemental**).

**Figure 1.**
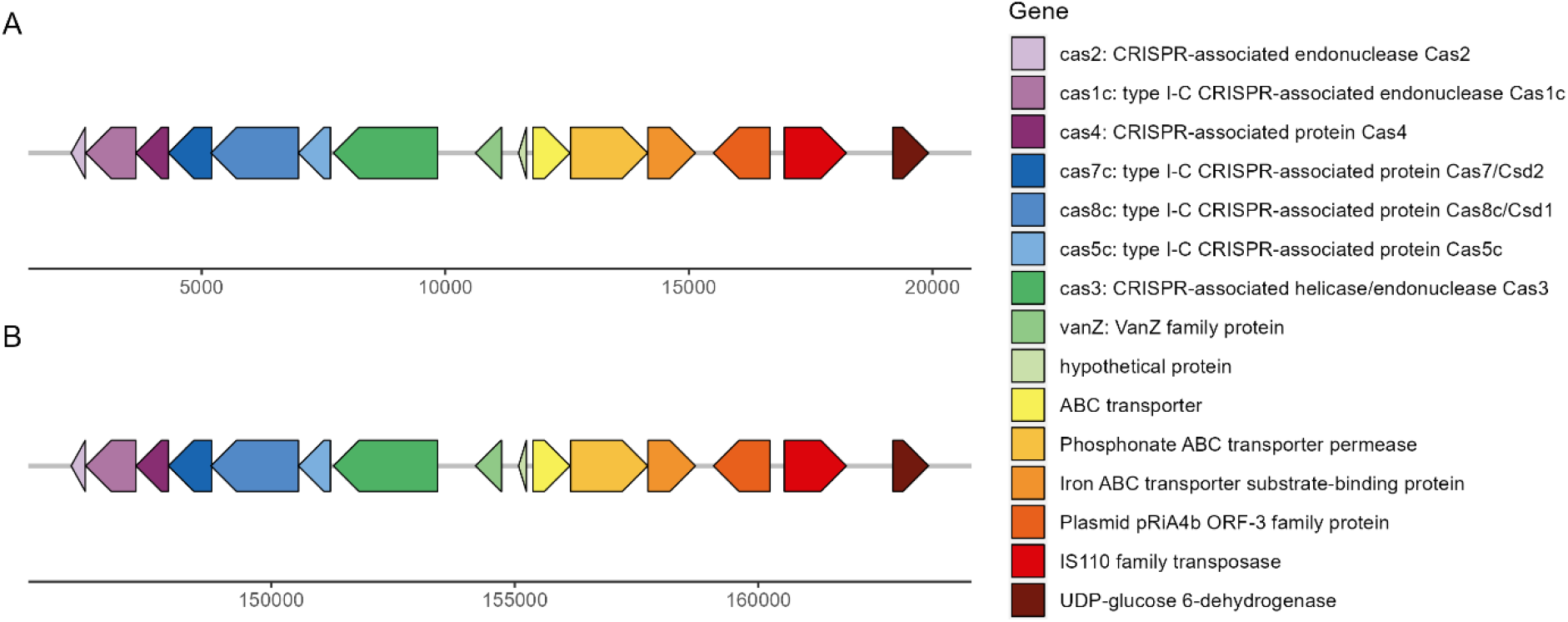
A VE202-06 genome region upstream and downstream of a gene encoding a VanZ family protein in A) the 2013 assembly and B) the 2022 WCB assembly. Workflow 2 gene annotations in a 17,598bp region of the **A)** 2013 assembly on contig BAHT02000202.1 or **B)** 2022 WCB assembly on the circular chromosome. Both genome regions contain identical gene annotations and synteny, as well as CRISPR-associated genes and a VanZ family protein.

Among the predicted gene products, VanR is the response regulator in the VanS-VanR two-component system, which transcriptionally activates glycopeptide resistance genes and thus is not sufficient to confer vancomycin resistance on its own^6^. VanY is not considered part of the core Van cascade and plays only an accessory role in resistance; expression of the *vanY* gene alone was not sufficient to confer vancomycin resistance in *E. faecalis*^6,13^. VanZ has an unknown role in the Van cascade, is dispensable in vancomycin resistance, and is not sufficient to confer resistance on its own in *Enterococcus*^6,14^. Indeed, genes annotated as VanZ are widespread across Gram-positive bacteria and are often found in typical human gut microbes (InterPro accession IPR006976). Members of the VanZ-superfamily proteins may provide a level of intrinsic resistance to glycopeptides through a reduction of binding of the antibiotics to the Gram-positive cell, through an unknown mechanism ^15^.

The results provided here show that the risk of VE202-06 serving as a donor of vancomycin resistance gene clusters to other members of the microbiota community is thus non-existent, as it lacks vancomycin resistance genes that are sufficient to confer resistance and there is no evidence that the predicted genes are mobile. We therefore conclude that there is no risk of vancomycin resistance transfer from strain VE202-06 to relevant microbial organisms in the recipient’s microbiota.

## AUTHOR CONTRIBUTIONS

J.M.N and A.R.W conceived the study. A.R.W and W.C performed the data analysis. W.V.S supported the interpretation of the results. J.M.N wrote the manuscript with input from all authors.

