## Supplemental figures for "Genomic Analyses Suggest No Risk of Vancomycin Resistance Transfer by Strain VE202-06"

**Supplementary Figure 1**

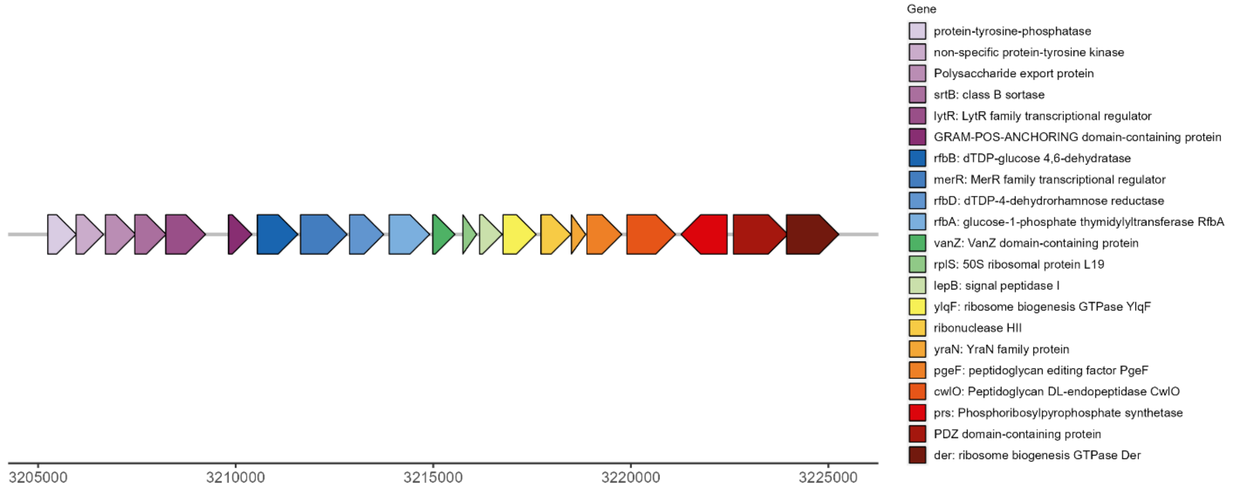


Workflow 2 gene annotations in a 20,561bp region of the 2022 WCB assembly surrounding a VanZ domain-containing protein gene.

**Supplementary Figure 2**

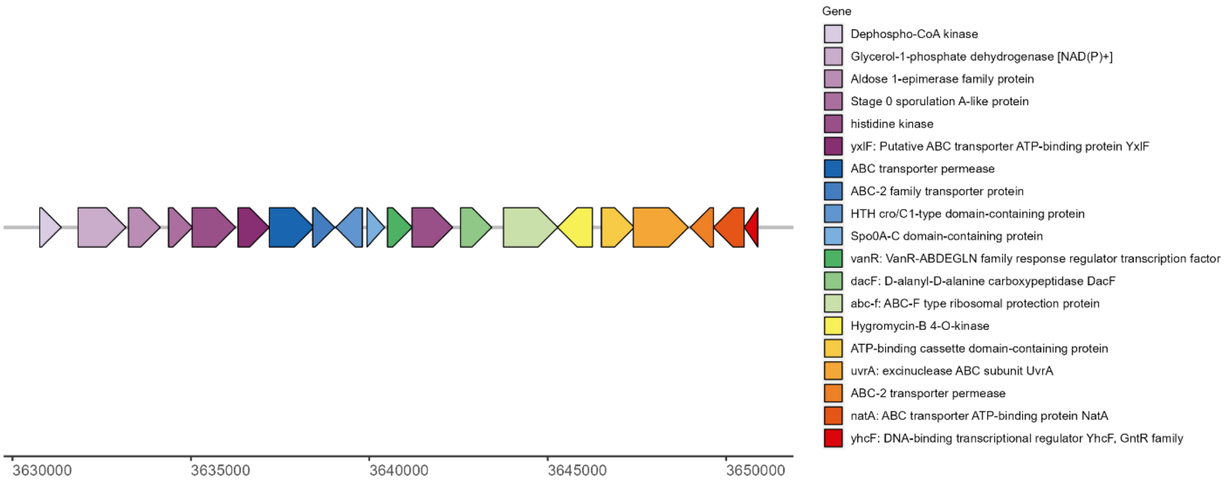


Workflow 2 gene annotations in a 20,696bp region of the 2022 WCB assembly surrounding a *vanR* gene.

**Supplementary Figure 3**

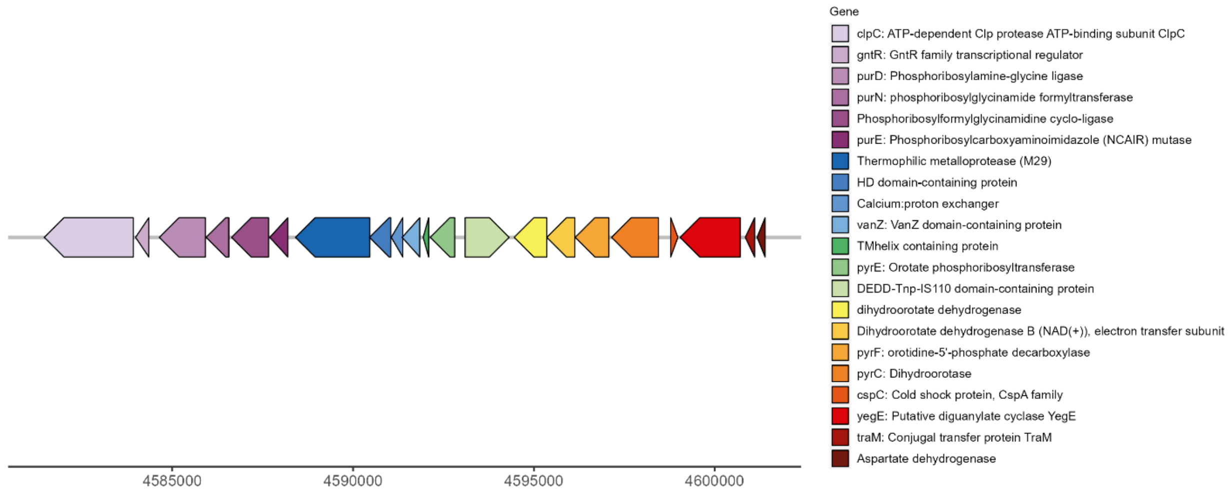


Workflow 2 gene annotations in a 20,489bp region of the 2022 WCB assembly surrounding a VanZ domain-containing protein gene.
